# Stressor- and tissue-specific regulation of the corticotropin-releasing factor system across epithelial tissues in rainbow trout

**DOI:** 10.64898/2026.06.11.731686

**Authors:** Brett M. Culbert, Alexis E. Pulford-Thorpe, Carol Best, Nicholas J. Bernier

## Abstract

The corticotropin-releasing factor (CRF) system is a major neural regulator of stress responses in vertebrates. However, stress-related roles for the CRF system in other tissues—and whether these roles vary between stressor types—remain unclear. To address this gap, we first characterized the CRF system in the gills and intestine of rainbow trout (*Oncorhynchus mykiss*) and then evaluated how it is transcriptionally regulated following either an immune (vaccination) or osmotic (seawater transfer) stressor. Additionally, since the CRF system is involved in food intake regulation, we also evaluated whether feeding state affects the intestinal CRF system. Vaccination against *Vibrio anguillarum* reduced CRF system activity in the intestine—as indicated by elevations in CRF binding protein transcripts paired with reductions in ligand (*crfa2*) and receptor (*crfr1b*) transcripts—but did not affect the gill CRF system. In contrast, seawater transfer caused the abundance of most CRF system transcripts to increase in the middle (but not posterior) portion of the intestine, while transcript levels of CRF binding proteins and receptors in the gills declined. Finally, levels of CRF system transcripts in the intestine varied with feeding state in a region-specific manner. In the middle intestine, transcript levels of most components declined with fasting and increased when feeding was resumed, whereas the opposite pattern occurred in the posterior intestine. Overall, our results implicate the peripheral CRF system as a stressor- and epithelial tissue-specific modulator of immune and osmoregulatory functions in teleosts.

## 1. Introduction

The ability to appropriately respond to environmental stressors is critical for an animal’s survival and fitness. To accomplish this, a suite of physiological responses occurs following a stressor (Carr, 2024; Wingfield, 2013), including elevated levels of two major classes of hormones: catecholamines (e.g., adrenaline and noradrenaline) and corticosteroids (e.g., cortisol and corticosterone). While catecholamines are critical for rapid physiological changes associated with ‘fight-or-flight’ responses [seconds to minutes; (Goldstein, 2025; Reid et al., 1998)], contributions of corticosteroids are generally slower and more prolonged (hours to days), facilitating acclimation to and/or recovery from a stressor (McEwen and Seeman, 1999). Given the relative ease with which corticosteroid levels can be measured and manipulated, a major focus of ecological and physiological studies alike has been to understand the regulation of and roles for these hormones during vertebrate stress responses (Bouyoucos et al., 2021; Creel et al., 2013; Dantzer et al., 2017). However, it is becoming increasingly clear that many hormone systems are necessary to overcome stressors and corticosteroids—while important—are but one of several hormones involved in coordinating stress responses (Chatzitomaris et al., 2017; Deck et al., 2017; Puglisi-Allegra and Andolina, 2015). Furthermore, the relative contribution of different hormone systems during stress responses varies depending on the nature, intensity, and duration of a stressor (Bernier et al., 2012, 2008; Bowers et al., 2008; Pacák and Palkovits, 2001). Yet, most studies continue to solely focus on the regulation of corticosteroids in response to environmental stressors and the relative contribution(s) of other stress-related hormone systems remain poorly understood.

The corticotropin-releasing factor (CRF) system—consisting of five ligands [CRFa, CRFb, urotensin 1 (UTS1), urocortin 2 (UCN2), and UCN3], a binding protein (CRFBP), and two receptors (CRFR1 and CRFR2) in teleost fishes (Culbert and Bernier, 2026; Maugars et al., 2022)—is the major neuroendocrine regulator of stress responses via its effects on the hypothalamic-pituitary-adrenal/interrenal axis and corticosteroid synthesis (Aguilera, 1998; Best et al., 2024). However, the CRF system also contributes to the regulation of many physiological processes outside of the brain. For example, activation of the CRF system stimulates proinflammatory responses in mammals (Chatoo et al., 2018; Dermitzaki et al., 2018; Kiank et al., 2010), counter to the anti-inflammatory effects that are generally associated with corticosteroids (Cain and Cidlowski, 2017). Similarly, the CRF system is an important regulator of osmoregulatory functions in the mammalian colon where it reduces ion and water absorption (Chatoo et al., 2018; Stengel and Taché, 2009). As in mammals, previous work in fish has suggested that the CRF system is involved in coordinating responses to immune challenges (Jiang et al., 2021; Mazon et al., 2006; Singh and Rai, 2011; van Heijningen et al., 2025) and osmotic disturbances (Aruna et al., 2021, 2012; Culbert et al., 2025a, 2025b, 2025c; Mainoya and Bern, 1982; Marshall and Bern, 1981). However, most of these studies have focused on only a small subset of CRF system components, and only Mazon et al. (2006) evaluated whether responses of the CRF system vary in a stressor- and tissue-specific manner using common carp (*Cyprinus carpio*). Therefore, we set out to compare whether transcriptional regulation of the CRF system varies between immune and osmotic stressors. We specifically focused on the gills and intestine because both tissues are key osmoregulatory organs (Evans et al., 2005; Grosell, 2010; Larsen et al., 2014) and are important for the prevention of pathogen entry from the external environment (Koppang et al., 2015; Salinas and Parra, 2015).

To accomplish this, we first characterized which components of the CRF system were most abundant in the gills and intestine of rainbow trout (*Oncorhynchus mykiss*). We then evaluated whether transcript levels of the major CRF system components in the gills and intestine were affected by either an immune (6, 24, 72, and 168h post-vaccination) or osmotic (24, 72, and 168h post-SW transfer) stressor. We predicted that vaccination would initially stimulate CRF system activity in the gill and intestine, associated with the initial production of proinflammatory cytokines, followed by a suppression of CRF system activity after production of anti-inflammatory cytokines increases. Following SW transfer, we predicted that activity of the gill CRF system would increase (reflecting the development of an ion-secretory phenotype) and intestine CRF system activity would decrease (reflecting a heightened need for ion and water absorption). Additionally, because the CRF system is a major regulator of food intake in teleosts (Bernier, 2006; Bernier and Peter, 2001; Conde-Sieira et al., 2018; Ortega et al., 2013) and salmonids naturally suppress feeding in response to stress—including osmotic and immune stressors (Craig et al., 2005; Sørum and Damsgård, 2004; Usher et al., 1991)—we also assessed whether feeding status influences transcript abundance of the intestinal CRF system. Specifically, we compared transcript abundance of CRF system components in the intestine of fed fish with fish that had been fasted for 0 (fed), 24, 72 or 168 h, as well as 24, 72, or 168 h after resumption of feeding following a 168 h fast. Here, we predicted that activity of the intestine CRF system would decline during fasting to maximize the capacity for nutrient absorption (i.e., reduced secretory capacity).

## 2. Methods

### 2.1. General Housing

All trout were acquired from the Ontario Aquaculture Research Centre (Alma, ON, Canada) and were housed in the Hagen Aqualab at the University of Guelph (Guelph, ON, Canada). Fish were initially maintained in 1.8m diameter fibreglass tanks (∼2000L) supplied with flow-through well water maintained at 12°C and kept on a 12h light, 12h dark photoperiod regime. Fish were fed to satiation three times per week with commercial trout food (3 or 5 PT Sinking, Blue Water Fish Food). A stocking density of ∼100 fish per tank was maintained during this period, and fish were kept under these conditions for at least a month prior to experiments. All procedures were carried out in accordance with the Canadian Council for Animal Care guidelines for the use of animals in research and teaching and were approved by the University of Guelph’s Animal Care Committee (AUPs #4123 and 4996).

### 2.2. Evaluating the CRF System of the Gills and Intestine

To determine which components of the CRF system were most abundant in the gills, as well as the middle (Mid. Int.) and posterior (Post. Int.) portions of the intestine, we sampled three trout (mass = 156 ± 10 g, fork length = 23.3 ± 0.9 cm) and performed qPCR (see Section 2.5) using gene-specific primers for each component (Supp. Table 1). All three fish were sexually immature.

**Table 1.** Time-dependent effects of vaccination (Exp 1), seawater transfer (Exp 2), and feeding state (Exp 3) on cortisol (ng mL^-1^; all Exp) and/or osmolality (mOsm kg^-1^; only Exp 2) in the plasma of rainbow trout (*Oncorhynchus mykiss*). Data are presented as means ± SEM (N = 7-11 per group). Significant effects are indicated with **bold** (p < 0.05). Differences as determined using post hoc analysis are depicted using either letters (time effects within a treatment, overall treatment effects, or overall time effects) or asterisks (treatment effect within a timepoint). Significant differences were not detected during post hoc analysis of plasma cortisol levels for Exp 3. Note that plasma cortisol and osmolality values from Exp 1 and 2 have previously been reported (Culbert et al., 2026 preprint and Culbert et al. 2025a).

| Variable | Exp | Treatment | Sampling Time |  |  |  |  | F | P |
| --- | --- | --- | --- | --- | --- | --- | --- | --- | --- |
| Plasma |  |  | 6h<sup>B</sup> | 24h<sup>A</sup> | 72h<sup>A</sup> | 168h<sup>A</sup> |  |  |  |
| Cortisol | 1 | Saline<sup>A</sup> | 7.4±1.6 | 3.1±0.9 | 1.1±0.3 | 4.6±0.7 | Treatment | 6.32 | 0.01 |
|  |  | Vaccine<sup>B</sup> | 11.3±1.3 | 4.4±1.0 | 3.2±0.8 | 3.3±0.5 | Time | 19.32 | <0.001 |
|  |  |  |  |  |  |  | Interaction | 2.63 | 0.06 |
|  |  |  |  | 24h | 72h | 168h |  |  |  |
|  | 2 | FW-to-FW |  | 12.9±3.5<sup>a</sup> | 6.4±3.2<sup>b</sup> | 10.4±2.7<sup>a</sup> | Treatment | 39.2 | <0.001 |
|  |  | FW-to-SW |  | 36.4±7.8<sup>AB*</sup> | 105.9±19.9<sup>B*</sup> | 14.9±4.5<sup>A</sup> | Time | 1.9 | 0.16 |
|  |  |  |  |  |  |  | Interaction | 12.3 | <0.001 |
|  |  |  | Fed | 24h | 72h | 168h |  |  |  |
|  | 3 | Fasted | 12.0±2.1 | 12.7±3.0 | 9.1±1.0 | 10.9±0.8 | Treatment | 3.71 | 0.005 |
|  |  | Fasted & Refed |  | 5.4±1.0 | 4.7±0.6 | 7.9±2.2 |  |  |  |
| Plasma |  |  |  | 24h | 72h | 168h |  |  |  |
| Osmolality | 2 | FW-to-FW<sup>A</sup> |  | 293±1 | 294±1 | 295±2 | Treatment | 177.8 | <0.001 |
|  |  | FW-to-SW<sup>B</sup> |  | 359±4 | 385±7 | 349±12 | Time | 2.8 | 0.07 |
|  |  |  |  |  |  |  | Interaction | 2.3 | 0.11 |

### 2.3. Experimental Design

#### 2.3.1. Experiment 1: Immune Challenge

Eighty-four sexually immature trout were used for this experiment (mass = 271 ± 6 g, fork length = 27.8 ± 0.2 cm). Fish were lightly anaesthetized using MS-222 (70 mg L^-1^), weighed, and subsequently transferred into one of eight experimental tanks (N=10-11 fish per tank). Fish were held in 0.6 m diameter fibreglass tanks (∼200L) containing an air stone. All tanks were supplied with flow-through well water maintained at 12°C and were kept on a 12h light, 12h dark photoperiod regime. Each tank was fed 1% of the average cumulative body weight in each tank daily and were held under these conditions for two weeks. Following this acclimation period, fish were anaesthetized using MS-222 (100 mg L^-1^) and injected intraperitoneally with 250 µL of either an inactivated vaccine solution or filter-sterilized saline (0.9% NaCl). Food was withheld for 24 h prior to injection. The vaccine (ICTHIOVAC VR, HIPRA, Spain) contained formalin-killed *Vibrio anguillarum* (serotypes O1, O2α and O2β) and has previously been shown to cause an immune response in rainbow trout (Khansari et al., 2019, 2018). After either 6, 24, 72, or 168 h post-injection, groups of vaccine- and saline-injected fish were killed via terminal anaesthesia using 2-phenoxyethanol (0.2%) at 1400 h. Fork length and mass were recorded, and blood was collected from the caudal vasculature using a 1 mL ammonium heparinized syringe. A portion of the blood was placed on ice to facilitate the evaluation of blood leukocyte abundance and respiratory burst activity (see Section 2.4.1), and the rest was centrifuged at 9,000 *g* for 5 min. After this, plasma was collected, frozen on dry ice, and stored at -80°C for later determination of cortisol levels (see Section 2.4.2). The intestinal tract was removed from each fish, flushed of its contents, and regionally dissected as previously described (Sundh et al., 2014). Specifically, we designated the middle intestine as the segment between the final pyloric caeca and the ileorectal sphincter, and the posterior intestine as the segment posterior to the ileorectal sphincter. These intestine regions, as well as the second gill arch on the right side of each fish, were frozen on dry ice and stored at -80°C to be used for later evaluation of immune-related transcripts and transcripts in the CRF system (see Section 2.5). Food was withheld throughout the experiment to avoid potential confounds associated with the inhibition of appetite due to vaccination (Sørum and Damsgård, 2004).

#### 2.3.2. Experiment 2: Osmotic Challenge

Fifty-eight trout were used for this experiment (mass = 544 ± 14 g, fork length = 35.0 ± 0.3 cm). Groups of ten fish were placed into recirculating tanks containing either freshwater or seawater (35ppt; Instant Ocean Sea Salt). All tanks contained ∼2000L of water at 12°C and were on a 12h light, 12h dark photoperiod regime. Additionally, all tanks were continuously aerated with an air stone and equipped with both particle and biological filtration, as well as UV sterilization. Each group of fish were sampled 24, 72, or 168 h following transfer. Two fish were removed from the 168 h FW-to-SW transferred group prior to the end of the experiment resulting in N = 8 for this group. Plasma (cortisol and osmolality; see Section 2.4.2), gill, and intestine segments (CRF system transcript abundance; see Section 2.5) were collected as described above. As in Experiment (Exp) 1, food was withheld throughout the experiment because salmonids naturally suppress food intake during seawater acclimation (Arnesen et al., 1993; Craig et al., 2005; Damsgård and Arnesen, 1998; Usher et al., 1991).

#### 2.3.3. Experiment 3: Effects of Fasting and Refeeding on the Intestine

Forty-nine trout were used for this experiment (mass = 750 ± 19 g, fork length = 37.9 ± 0.3 cm; N = 27 females and 22 males) and most (>80%) were still juvenile. Fish were lightly anaesthetized using MS-222 (70 mg L^-1^), weighed, and subsequently transferred into the experimental tanks (N=7 fish per tank). Fish were held in 1.2 m diameter fibreglass tanks (∼500L) that were supplied with aerated, flow-through well water. Tanks were maintained at 12°C and kept on a 12h light, 12h dark photoperiod regime. For two weeks prior to the start of fasting, fish were fed 1.5% of the average cumulative body weight in each tank at 1100 h every day (a pilot study determined this to be the maximum amount that fish would fully consume daily). Following this acclimation period, all fish in one tank were immediately sacrificed (Control). Fish in the other six tanks were sampled following either 24, 72, or 168 h of fasting, or 24, 72, or 168 h after feeding had resumed following a 168 h fast. All treatment groups were terminally anaesthetized at 1200 h (1 h after feeding for non-fasted groups), and plasma (cortisol; see Section 2.4.2) and intestine (CRF system transcript abundance; see Section 2.5) were collected as described above.

### 2.4. Physiological Parameters

#### 2.4.1. Immune Indicators

For Exp 1, we evaluated several immune-related variables to confirm that our vaccination protocol was effective in eliciting an immune response at both whole animal (leukocyte counts and respiratory burst in the blood; see below) and tissue levels (transcript abundance of immune-related genes in each tissue of interest; see section 2.5).

The total proportion of circulating leukocytes—as well as separate subpopulations of lymphocytes/thrombocytes (called ‘lymphocytes’) and neutrophil/monocytes (called ‘neutrophils’)—was quantified using flow cytometry according to the protocol of Inoue et al. (2002). Briefly, 10 µL of whole blood was diluted into 1986 µL of Hank’s balanced salt solution (HBSS; Item #H8264, Sigma-Aldrich) and 4 µL of ethanol containing 500 µg mL^-1^ DiOC_6_ (3,3′-dihexyloxacarbocyanine iodide; Item #318426, Sigma-Aldrich). After incubating at room temperature for 10 mins in the dark with shaking (100 rpm), samples were filtered with a cell strainer (35 µm, Falcon) and loaded into a FACSMelody cell sorter (BD Biosciences, RRID:SCR_023209). All samples were agitated at 200 rpm during analysis, and an event rate of ∼6000 events s^-1^ was maintained until 1 million events were recorded. Voltage thresholds were set at 15, 49, and 32 volts for forward scatter, side scatter, and fluorescence photomultiplier tubes, respectively. Leukocyte subpopulations (see above) were identified within the FACSChorus software (BD Biosciences) based on differences in forward scatter, side scatter, and fluorescence intensities, as previously described (Inoue et al., 2002).

The respiratory burst activity of leukocytes in the blood was measured as previously described (Semple et al., 2018). Briefly, white polystyrene microplates were prepared with four wells containing 75 µL of whole blood from each fish (three stimulated and one unstimulated). Stimulated wells received 85 µL of HBSS, 40 µL of luminol in HBSS (10 mM; Item #A8511, Sigma-Aldrich), and 100 µL of zymosan A in HBSS (20 mg mL^-1^; Item #Z4250, Sigma-Aldrich). Unstimulated wells received the same reagents except that 100 µL of HBSS was added in lieu of zymosan solution. The chemiluminescent signal in each well was measured every 3 min over 150 min using an LMax II^384^ luminometer (Molecular Devices). We recorded the integral of the signal over the entire duration and, to account for background signal, the value of the unstimulated well was subtracted from the average value of the three stimulated wells for each fish (intra-assay variation was 8.8% CV).

#### 2.4.2. Plasma Cortisol and Osmolality

Plasma cortisol levels were measured using a previously validated, commercially available enzyme-linked immunosorbent assay (ELISA; Item #402710, Neogen) for fish from all experiments. Values of intra- and inter-assay variation were 9.3% and 4.5% CV, respectively. Additionally, plasma osmolality values were determined in duplicate using a vapour pressure osmometer (Vapro 5520, Wescor) for fish from Exp 2, which had an intra-assay variation of 0.8% CV.

### 2.5. RNA Isolation and qPCR

Gill and intestine (both middle and posterior regions) samples were ground on dry ice with a mortar and pestle, then homogenized in Ribozol reagent (VWR International) using a Precellys Evolution tissue homogenizer (Bertin Instruments). Total RNA was extracted following the manufacturer’s protocol, after which its quantity and purity were assessed using a NanoDrop 2000 spectrophotometer (Thermo Scientific). We then treated 1 µg of RNA with DNase (DNase 1; Thermo Scientific) and reverse-transcribed cDNA using a high-capacity cDNA reverse transcription kit (Applied Biosystems). We performed qPCR using a CFX96 system (BioRad, RRID:SCR_018064) with SYBR green (SsoAdvanced Universal; BioRad) and gene-specific primers (Supp. Table 1). All samples were run in duplicate, and negative controls, including no-template controls (where cDNA was replaced with water) and no reverse transcriptase controls (where RNA reverse transcriptase was replaced with water during cDNA synthesis) were included. Each reaction contained a total of 20 µl, which consisted of 10 µl of SYBR green, 5 µl of combined forward and reverse primers (0.2 µM [final]), and 5 µl of 10x diluted cDNA. Cycling parameters included a 30 s activation step at 95°C, followed by 40 cycles consisting of a 3 s denaturation step at 95°C and a combined 30 s annealing and extension step at 60°C. Melt curve analysis was conducted at the end of each run to confirm the specificity of each reaction. To account for differences in amplification efficiency, standard curves were constructed for each gene using serial dilutions (4x) of pooled cDNA. Input values for each gene were obtained by fitting the average threshold cycle value to the antilog of the gene-specific standard curve, thereby correcting for differences in primer amplification efficiency. To correct for minor variations in template input and transcriptional efficiency, we measured transcript abundance of elongation factor 1 α (*ef1α*) and ribosomal protein L13a (*rpl13a*) as reference genes, which were both stable across groups in all experiments. Data were normalized using the geometric mean of these reference genes and are expressed relative to the mean value of the 6h saline group in Exp 1, the 24h FW group in Exp 2, and the fed group in Exp 3.

### 2.6. Statistical Analysis

Statistical analyses were performed using R (version 4.4.0, RRID:SCR_001905). All data are presented as means ± 1 standard error of the mean (SEM), and a significance level (α) of 0.05 was used for all tests. Outliers were excluded based on a 2× interquartile range threshold. When data did not meet the assumptions of normality and/or equal variance, data were transformed to improve model fit. Data from each time course series was analyzed using ANOVAs that included either only time as a factor (Exp 3) or time, treatment, and their interaction term as factors (Exp 1 and 2). All models were fit using the lm function in the ’lme4’ package [RRID:SCR_015654 (Bates et al., 2015)] and overall differences were determined using the Anova function in the ‘car’ package (RRID:SCR_022137). When significant differences were detected, Tukey’s HSD post hoc analysis was performed using the emmeans function in the ‘emmeans’ package [RRID:SCR_018734 (Lenth, 2016)]. Sex was not included in our analyses because fish were either sexually immature (Exps 1 and 2) or preliminary analyses revealed no effects of sex (Exp 3). The major findings of our analyses are described in the results section below; however, a detailed description of all non-significant statistical results can be found in the electronic supplementary material (Supp. Tables 2-4).

**Table 2.**
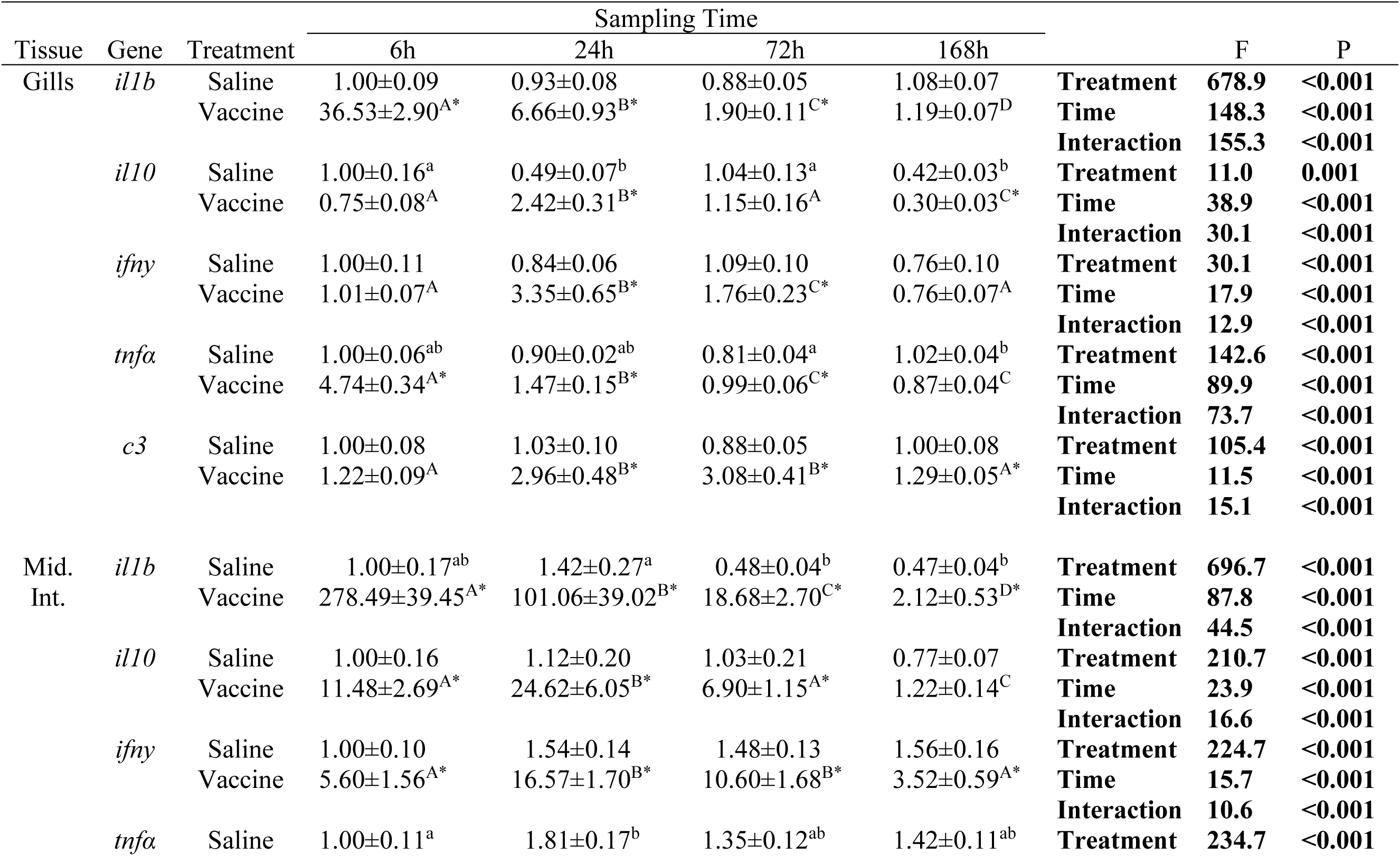
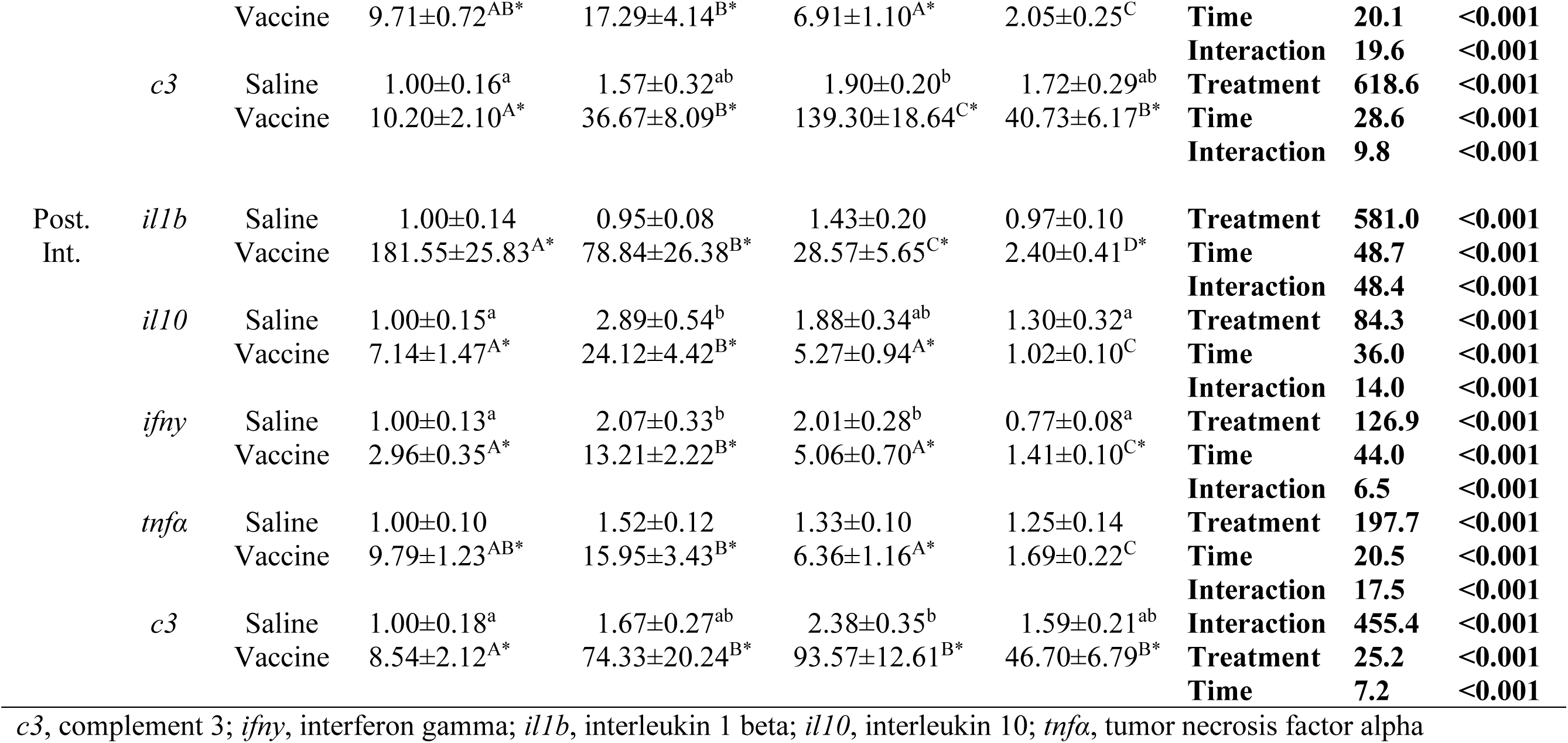
Time-dependent effects of saline or vaccine treatment on immune-related genes in the gills, middle intestine (Mid. Int.), and posterior intestine (Post. Int.) of rainbow trout (*Oncorhynchus mykiss*). Data are presented as means ± SEM (N = 8-11 per group) and are expressed relative to the saline group at 6h. Significant effects are indicated with **bold** (p < 0.05). Differences as determined using post hoc analysis are depicted using either letters (time effects within a treatment) or asterisks (treatment effect within a timepoint).

## 3. Results

### Tissue Distribution

In general, transcript abundance of binding proteins and receptors was far more abundant in the gills and both regions of the intestine compared to most ligands (Fig. 1). While undetectable in either region of the intestine, *crfr2a* was the most abundant receptor in the gills. Similarly, *crfr1a* and *crfr1b* were both ∼5x more abundant in the gills than in either the Mid. Int. or Post. Int. Levels of *crfr2b* were twice as high in the Post. Int. compared to levels in either the gills or Mid. Int. For binding proteins, *crfbp1* was the most abundant component expressed in the gills and was ∼30x and ∼60x more abundant there than in the Mid. Int. and Post. Int., respectively. In contrast, *crfbp2* was the most abundant component in the intestine, with ∼60x and ∼30x greater abundance in the Mid. Int. and Post. Int., respectively, compared to the gills. Lastly, *ucn3* was ∼4x more abundant in the gills than in the intestine, *crfa2* was ∼5x more abundant in the intestine than in the gills, and *crfa1* levels were low but detectable in all tissues. In contrast, we only detected *ucn2b* in the intestine, *crfb1* and *uts1a* in the gills, and none of *crfb2*, *uts1b*, or *ucn2a* were detectable in any of the examined tissues.

**Figure 1.**
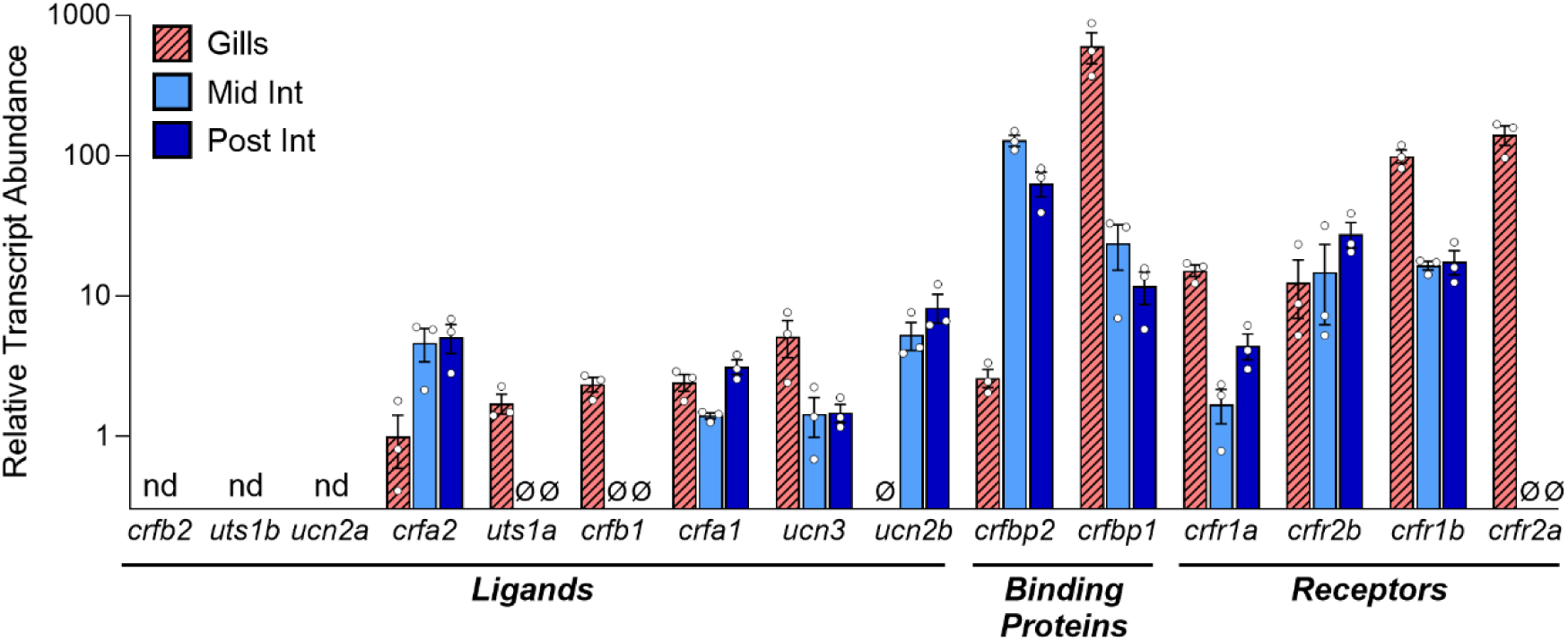
Relative abundance of individual ligands, binding proteins, and receptors of the corticotropin-releasing factor system in the gills, middle intestine (Mid. Int.), and posterior intestine (Post. Int.) of rainbow trout (*Oncorhynchus mykiss*). Bars represent mean abundance (±SEM) of each component relative to *crfa2* in the gills (the component with the lowest detectable levels). Each point (N=3) represents an individual value from an unstressed fish. Components that were not detected in any of the three tissues are indicated with nd (not detected), and the Ø symbol indicates instances where components were only undetectable in some tissues. Note that data are plotted on a Log10 scale for visualization purposes. *crf*, corticotropin-releasing factor; *crfbp*, corticotropin-releasing factor binding protein; *crfr*, corticotropin-releasing factor receptor; *ucn*, urocortin; *uts*, urotensin.

### Experiment 1: Immune challenge

Vaccination caused total circulating leukocytes to decrease by 50% at 24h (Fig. 2A; p_treatment_=0.63, p_time_<0.001, p_interaction_<0.001), which was driven by a concurrent 50% reduction in lymphocyte counts at 24h (Fig. 2B; p_treatment_=0.02, p_time_<0.001, p_interaction_<0.001). In contrast, blood neutrophil counts in vaccinated fish were ∼2.3 and ∼1.5x higher at 72h and 168h (Fig. 2C; p_treatment_<0.001, p_time_<0.001, p_interaction_<0.001), respectively. Similarly, respiratory burst activity in the blood was ∼2.5x and ∼1.5x greater in vaccinated fish at 72h and 168h (Fig. 2D; p_treatment_=0.03, p_time_<0.001, p_interaction_=0.04), respectively. Plasma cortisol levels were ∼35% higher in vaccinated fish across all time points and ∼3x higher at 6h compared to all other time points in both groups (Table 1); however, plasma cortisol levels were generally low in all fish (<10 ng mL^-1^). Consistent with the observed immunological responses in the blood, transcript abundance of immune-related genes further confirmed that all examined tissues mounted a robust immune response (Table 2). Peak activation of proinflammatory cytokines (*il1b*, *ifnγ*, and *tnfα* at 6-24h) was followed by the elevation of a key anti-inflammatory cytokine (*il10* at 24h), as well as a delayed and prolonged activation of the complement system (*c3* at 24-168h). Transcriptional responses of all immune-related genes were greater in both regions of the intestine compared to the gills.

**Figure 2.**
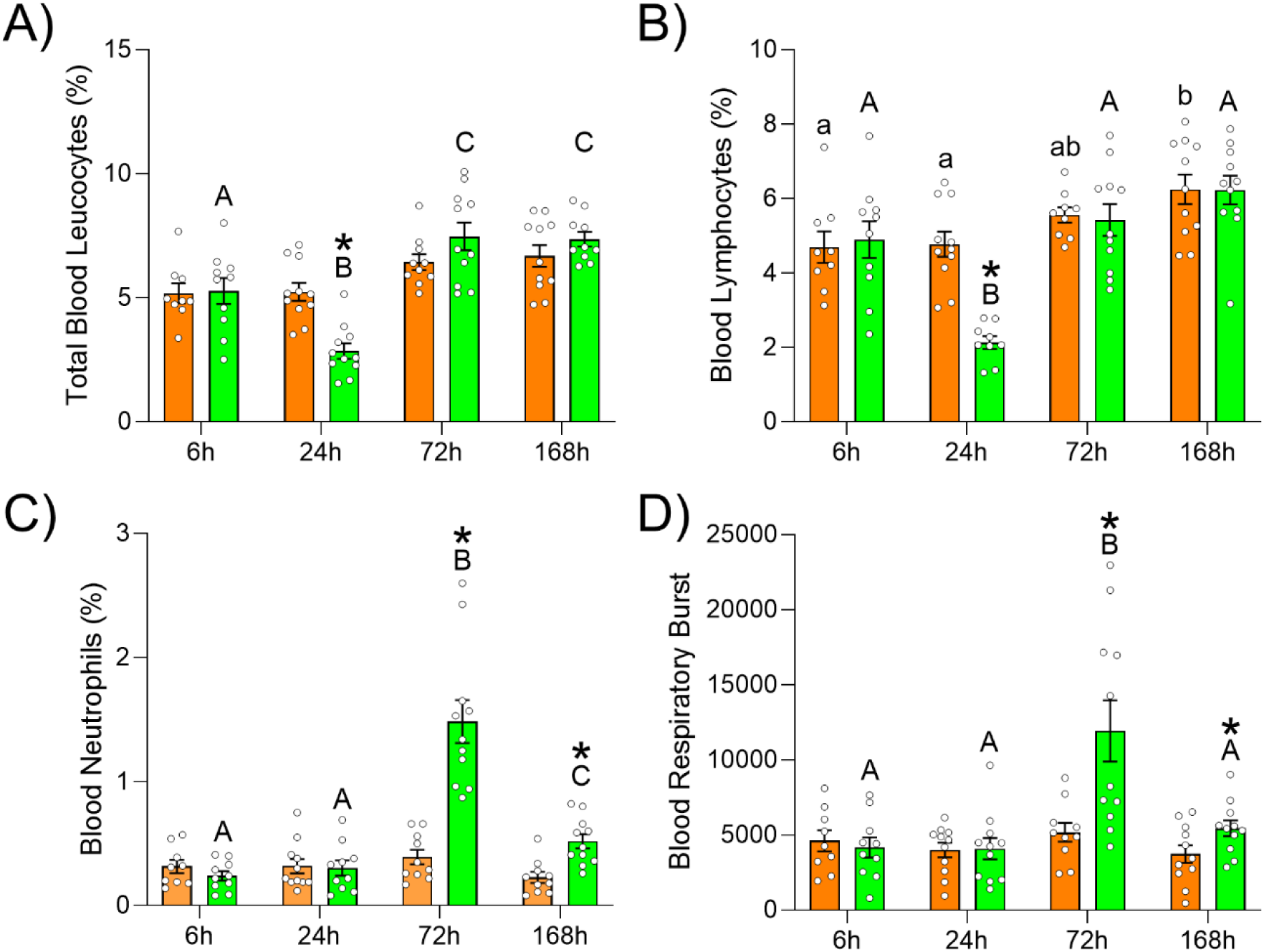
Time-dependent effects of saline or vaccine treatment on the percentage of leukocytes (A), lymphocytes (B), and neutrophils (C), as well as respiratory burst (D) in the blood of rainbow trout (*Oncorhynchus mykiss*). Significant differences (p < 0.05; two-way ANOVA) are depicted using either letters (across time; uppercase = within vaccine group, lowercase = within saline group) or asterisks (between groups within a timepoint). Values are represented as means ± SEM, and individual data points are shown (N=9-11).

In the Mid. Int., levels of *crfbp1* (Fig. 3A) and *crfbp2* (Fig. 3B) were both elevated in vaccinated fish compared to saline-treated controls. Specifically, *crfbp1* was ∼2.5x and ∼2x higher at 24 and 72h (p_treatment_<0.001, p_time_<0.001, p_interaction_<0.001), respectively, while *crfbp2* in vaccinated fish was ∼25% higher overall (p_treatment_=0.04, p_time_=0.41, p_interaction_=0.32). In contrast, levels of *crfr1b* (Fig. 3C; p_treatment_=0.03, p_time_<0.001, p_interaction_=0.19) and *crfa2* (Fig. 3D; p_treatment_=0.002, p_time_<0.001, p_interaction_=0.59) were lower in vaccinated fish compared to saline-treated fish. This effect was greatest at 24h for *crfr1b*—with levels in vaccinated fish being 25% those of saline-treated fish—but levels of *crfa2* were ∼20% lower in vaccinated fish across all time points. Additionally, the abundance of *crfr1b* and *crfa2* was 2-3-fold higher overall at 24, 72, and 168h compared to 6h. No vaccine-related changes were observed for *crfr2b* or *ucn2b* (Supp. Table 2).

**Figure 3.**
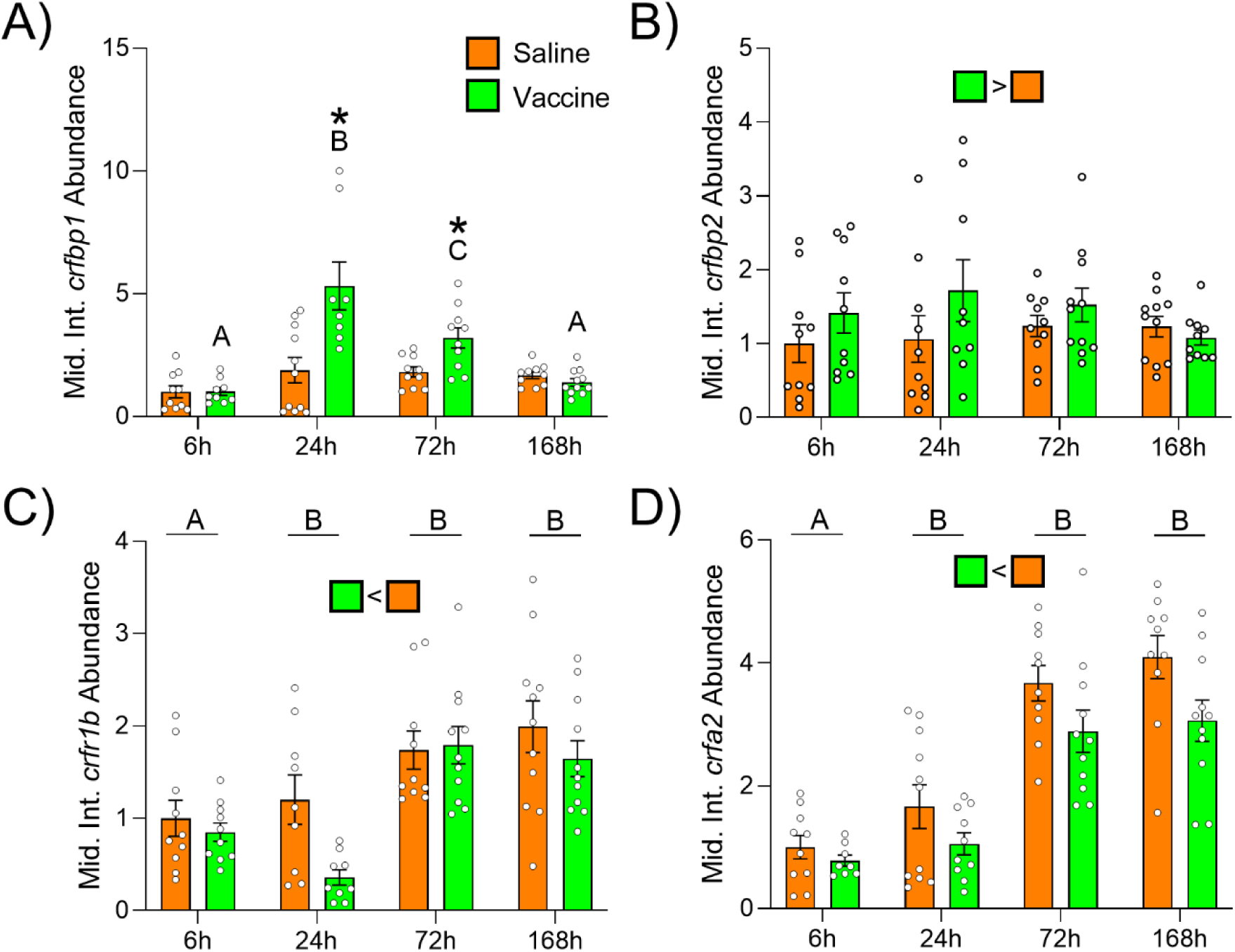
Time-dependent effects of saline or vaccine treatment on transcript abundance of corticotropin-releasing factor (CRF) binding protein 1 (*crfbp1*; A), CRF binding protein 2 (*crfbp2*; B), CRF receptor 1b (*crfr1b*; C), and CRFa2 (*crfa2*; D) in the middle intestine (Mid. Int.) of rainbow trout (*Oncorhynchus mykiss*). Significant differences (p < 0.05; two-way ANOVA) are depicted using either letters (across time; uppercase = within vaccine group, underlined uppercase = overall time effect), filled oversized squares (between groups across all timepoints) or asterisks (between groups within a timepoint). Data are expressed relative to saline-injected fish at 6h. Values are represented as means ± SEM, and individual data points are shown (N=8-11).

In the Post. Int., changes in *crfbp1* (Fig. 4A) and *crfr1b* (Fig. 4B) mirrored those observed in the Mid. Int. Specifically, levels of *crfbp1* in vaccinated fish were ∼2.5x higher than those of saline-treated fish at both 24 and 72h (p_treatment_<0.001, p_time_<0.001, p_interaction_=0.006), and levels of *crfr1b* were ∼70% lower than those of saline-treated fish at 24h (p_treatment_=0.08, p_time_=0.21, p_interaction_=0.01). While unaffected by vaccination, levels of *crfa2* (Fig. 4C; p_treatment_=0.62, p_time_<0.001, p_interaction_=0.07) gradually increased across the duration of the experiment with levels at 168h being twice those at 6h. No vaccine-related changes were observed for *crfbp2*, *crfr2b*, or *ucn2b* (Supp. Table 2).

**Figure 4.**
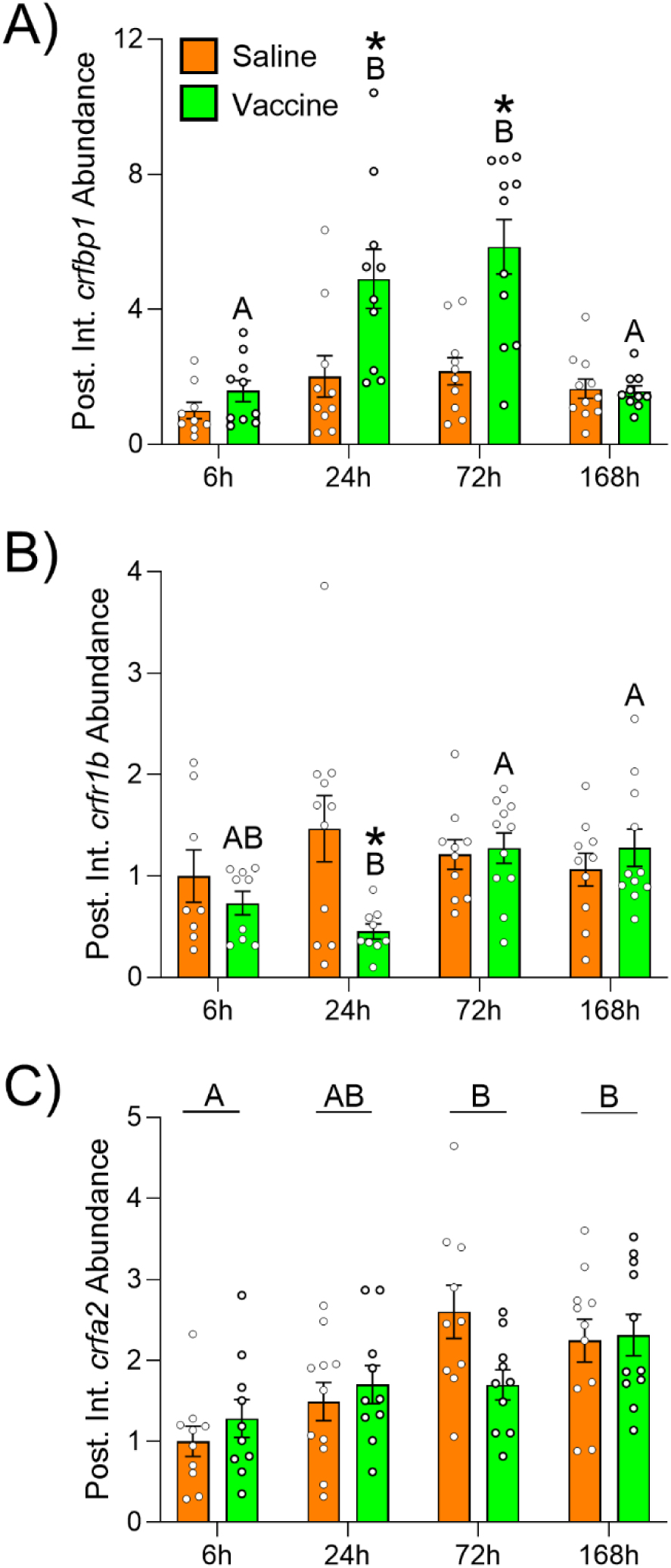
Time-dependent effects of saline or vaccine treatment on transcript abundance of corticotropin-releasing factor (CRF) binding protein 1 (*crfbp1*; A), CRF receptor 1b (*crfr1b*; B), and CRFa2 (*crfa2*; C) in the posterior intestine (Post. Int.) of rainbow trout (*Oncorhynchus mykiss*). Significant differences (p < 0.05; two-way ANOVA) are depicted using either letters (across time; uppercase = within vaccine group, underlined uppercase = overall time effect) or asterisks (between groups within a timepoint). Data are expressed relative to saline-injected fish at 6h. Values are represented as means ± SEM, and individual data points are shown (N=8-11).

In contrast to the intestine, the only vaccine-related change observed in the gills was lower *crfr2a* levels in vaccinated fish compared to saline-treated fish at 72h (Supp. Table 2). However, this difference was primarily driven by a transient reduction in the saline group at 72h and not by altered *crfr2a* levels in vaccinated fish. Similarly, no vaccine-related changes were observed for *crfbp1*, *crfr1a*, *crfr1b*, *crfr2b*, or *ucn3* (Supp. Table 2).

### Experiment 2: Osmotic challenge

As previously reported (Culbert et al., 2025a), plasma cortisol and osmolality values (Table 1) were both elevated in SW versus FW fish. Plasma cortisol levels were ∼3x and ∼17x higher in SW fish at 24 and 72h, respectively, whereas osmolality was ∼25% higher in SW fish across the duration of the experiment.

In the Mid. Int., time-dependent increases in all CRF components were observed in SW fish. Levels of *crfbp1* (Fig. 5A) transiently increased 2.5-fold in SW fish at 72h (p_treatment_=0.04, p_time_=0.86, p_interaction_<0.001), whereas levels of *crfbp2* (Fig. 5B; p_treatment_=0.70, p_time_<0.001, p_interaction_=0.001), *crfr1b* (Fig. 5C; p_treatment_=0.20, p_time_<0.001, p_interaction_<0.001) and *crfr2b* (Fig. 5D; p_treatment_=0.08, p_time_<0.001, p_interaction_<0.001) in SW fish were ∼50% those of FW fish at 24h, but had increased to levels ∼2-fold higher than FW fish by 168h. Levels of *crfa2* (Fig. 5E; p_treatment_=0.20, p_time_<0.001, p_interaction_<0.001) and *ucn2b* (Fig. 5F; p_treatment_=0.03, p_time_=0.003, p_interaction_=0.32) were also higher in SW fish compared to FW fish—especially at 72 (∼3x higher) and 168h (∼60% higher). Additionally, transcript levels of both peptides increased 2-3-fold over the course of the experiment.

**Figure 5.**
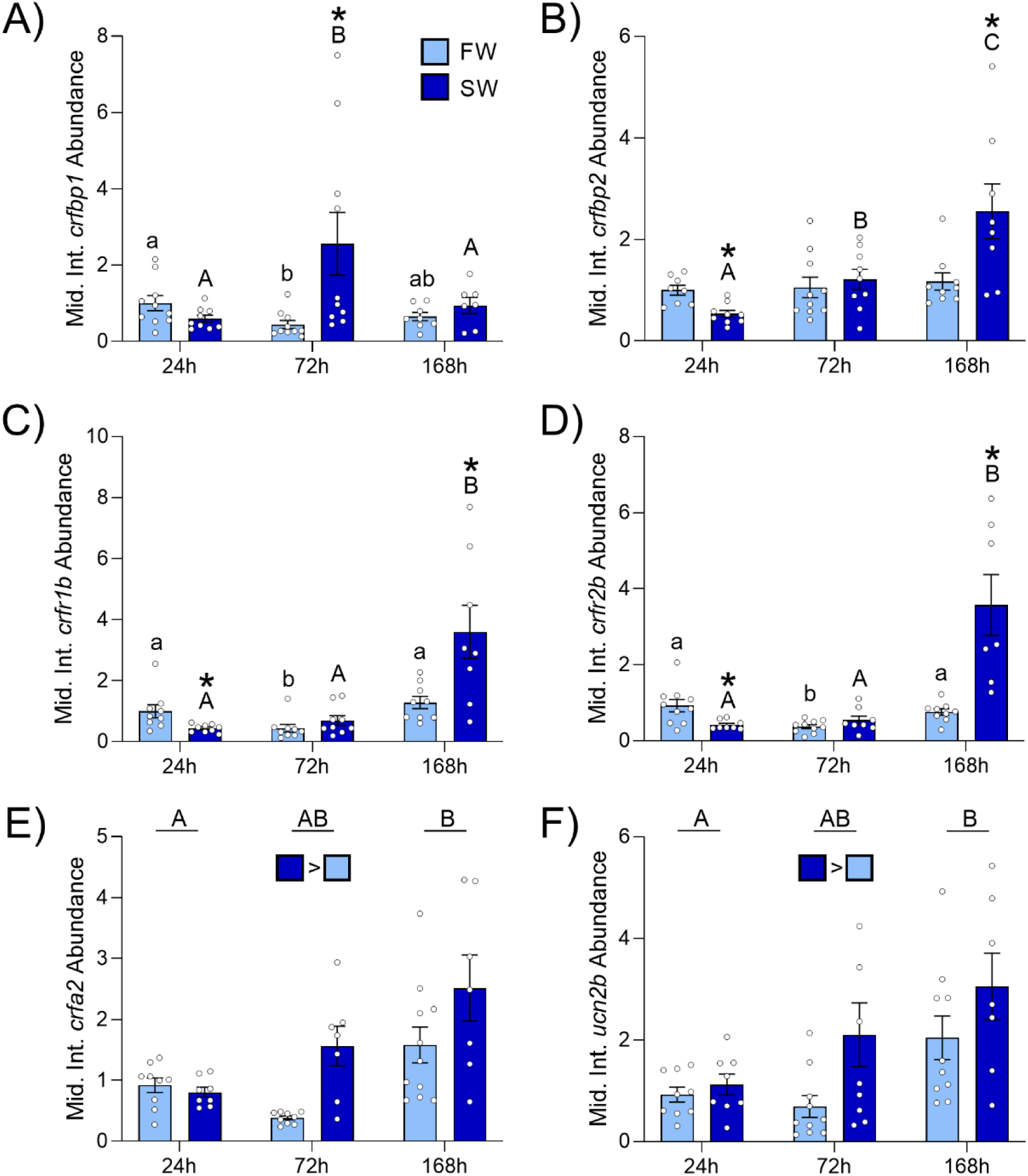
Time-dependent effects of transfer from freshwater-to-freshwater (FW) or freshwater-to-seawater (SW) on transcript abundance of corticotropin-releasing factor (CRF) binding protein 1 (*crfbp1*; A), CRF binding protein 2 (*crfbp2*; B), CRF receptors 1b (*crfr1b*; C) and 2b (*crfr2b*; D), CRFa2 (*crfa2*; E), and urocortin 2b (*ucn2b*; F) in the middle intestine (Mid. Int.) of rainbow trout (*Oncorhynchus mykiss*). Significant differences (p < 0.05; two-way ANOVA) are depicted using either letters (across time; uppercase = within SW, lowercase = within FW, underlined uppercase = overall time effect), filled oversized squares (between groups across all timepoints) or asterisks (between groups within a timepoint). Data are expressed relative to FW fish at 24h. Values are represented as means ± SEM, and individual data points are shown (N=7-10).

In the Post. Int., levels of *crfbp1* (Fig. 6A) increased ∼3-fold across time in FW fish but not in SW fish (p_treatment_=0.27, p_time_=0.10, p_interaction_=0.046). Similar increases of ∼2-3-fold across the duration of the experiment were observed for *crfbp2* (Fig. 6B; p_treatment_=0.85, p_time_=0.03, p_interaction_=0.68), *crfr1b* (Fig. 6C; p_treatment_=0.71, p_time_=0.06, p_interaction_=0.99), *crfa2* (Fig. 6D; p_treatment_=0.48, p_time_=0.04, p_interaction_=0.08), and *ucn2b* (Fig. 6E; p_treatment_=0.97, p_time_=0.03, p_interaction_=0.85); however, these changes were consistent between FW and SW fish. No differences were detected for *crfr2b* (Supp. Table 3).

**Figure 6.**
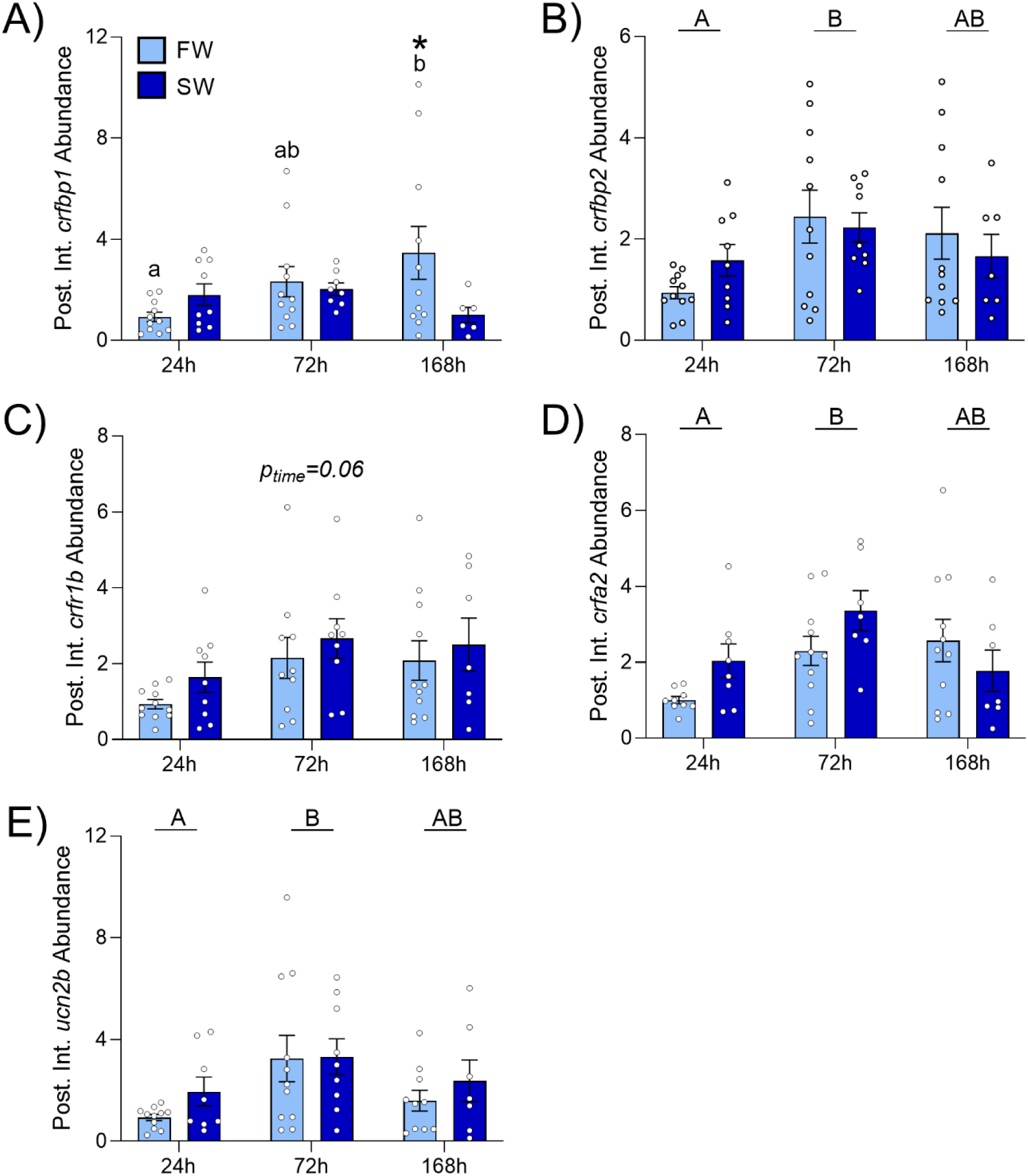
Time-dependent effects of transfer from freshwater-to-freshwater (FW) or freshwater-to-seawater (SW) on transcript abundance of corticotropin-releasing factor (CRF) binding proteins 1 (*crfbp1*; A) and 2 (*crfbp2*; B), CRF receptor 1b (*crfr1b*; C), CRFa2 (*crfa2*; D), and urocortin 2b (*ucn2b*; E) in the posterior intestine (Post. Int.) of rainbow trout (*Oncorhynchus mykiss*). Significant differences (p < 0.05; two-way ANOVA) are depicted using either letters (across time; lowercase = within FW, underlined uppercase = overall time effect) or asterisks (between groups within a timepoint). Data are expressed relative to FW fish at 24h. Values are represented as means ± SEM and individual data points are shown (N=7-10).

In the gills, levels of *crfbp1* (Fig. 7A), *crfr1a* (Fig. 7B), *crfr1b* (Fig. 7C), and *crfr2a* (Fig. 7D) were all lower in SW fish. Time-dependent reductions of ∼30% were detected in SW fish for *crfbp1* (p_treatment_=0.44, p_time_=0.02, p_interaction_=0.008) and *crfr1b* (p_treatment_=0.10, p_time_=0.74, p_interaction_=0.04) at 168h and 72h, respectively. Levels of *crfr1a* (p_treatment_=0.003, p_time_=0.65, p_interaction_=0.77) and *crfr2a* (p_treatment_<0.001, p_time_=0.46, p_interaction_=0.20) in SW fish were ∼30% and ∼40% lower, respectively, across all timepoints. No differences in *crfr2b* or *ucn3* were detected (Supp. Table 3).

**Figure 7.**
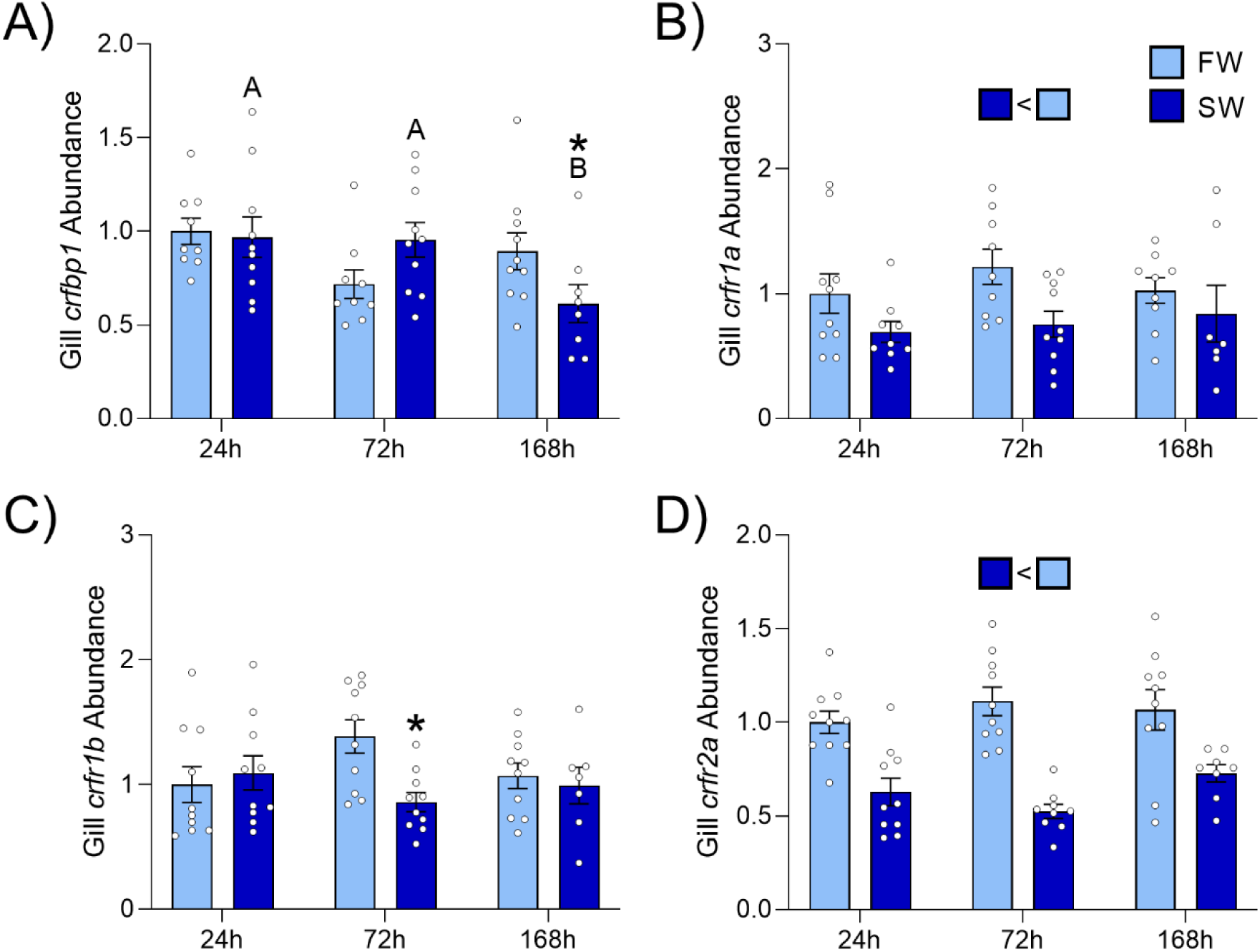
Time-dependent effects of transfer from freshwater-to-freshwater (FW) or freshwater-to-seawater (SW) on transcript abundance of corticotropin-releasing factor (CRF) binding protein 1 (*crfbp1*; A), and CRF receptors 1a (*crfr1a*; B), 1b (*crfr1b*; C), and 2a (*crfr2a*; D) in the gills of rainbow trout (*Oncorhynchus mykiss*). Significant differences (p < 0.05; two-way ANOVA) are depicted using either letters (across time within SW group), filled oversized squares (between groups across all timepoints) or asterisks (between groups within a timepoint). Data are expressed relative to FW fish at 24h. Values are represented as means ± SEM and individual data points are shown (N=7-10).

### Experiment 3: Fasting and re-feeding

In the Mid. Int., abundance of CRF system components—specifically, *crfbp1* (Fig. 8A; p=0.03), *crfbp2* (Fig. 8B; p=0.03), *crfr1b* (Fig. 8C; p=0.01), and *crfr2b* (Fig. 8D; p<0.001)—generally decreased (by up to 50%) over the course of the 7d fast. Transcript levels gradually returned to levels consistent with fed animals once feeding resumed. In the Post. Int., levels of *crfbp2* (Fig. 9A; p<0.001), *crfr1b* (Fig. 9B; p=0.009), *crfr2b* (Fig. 9C; p=0.02), and *crfa2* (Fig. 9D; p=0.02) all increased ∼2-4-fold during the 7d fast. Interestingly, while abundance of these genes all returned to levels consistent with fed animals by the end of the 7d re-feeding period, levels of *crfbp2*, *crfr1b*, and *crfr2b* initially increased ∼2-3-fold prior to this decline. No changes were observed in *crfa2* and *ucn2b* in the Mid. Int. or *ucn2b* and *crfbp1* in the Post. Int. (Supp. Table 4).

**Figure 8.**
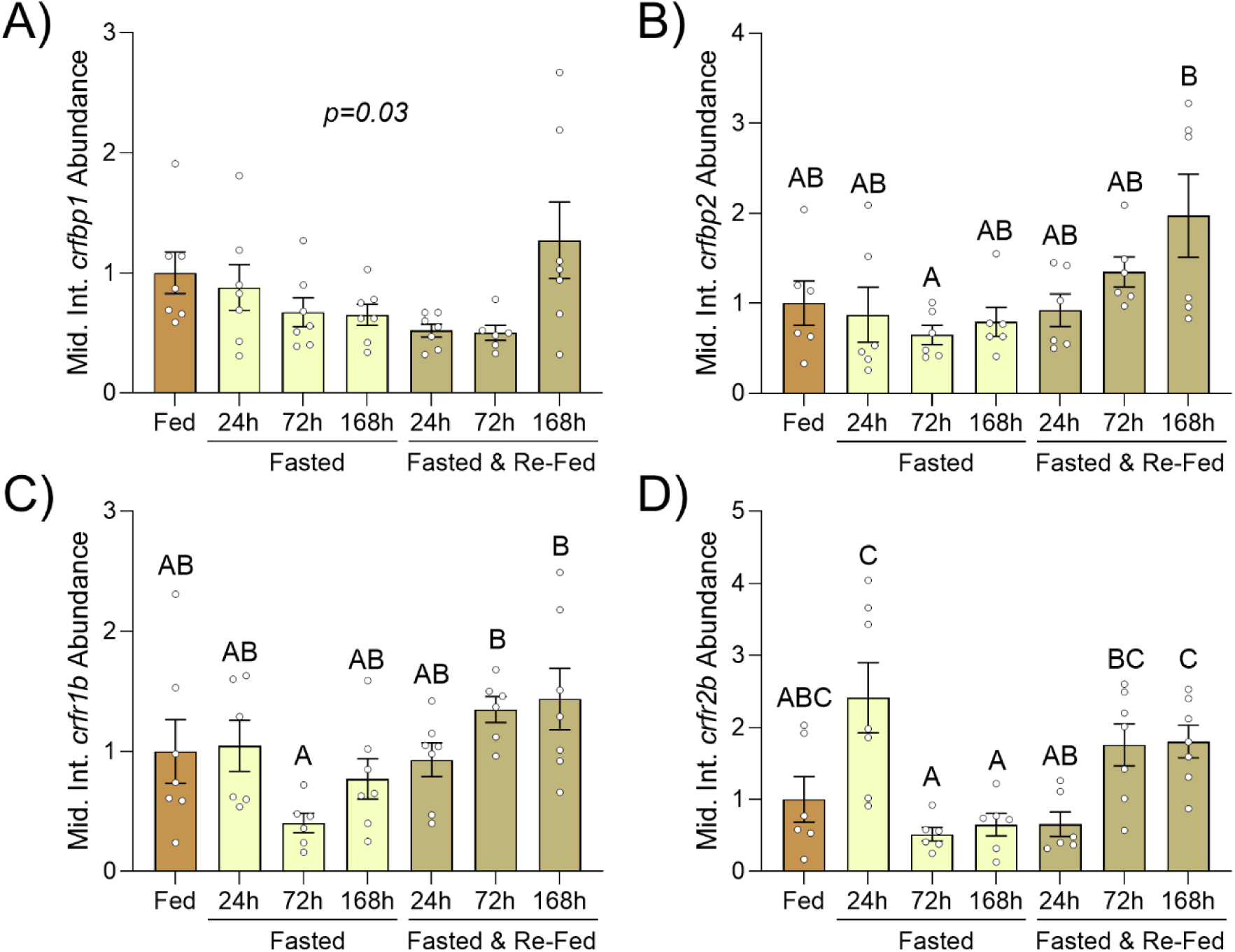
Time-dependent effects of fasting and re-feeding on transcript abundance of corticotropin-releasing factor (CRF) binding proteins 1 (*crfbp1*; A) and 2 (*crfbp2*; B), and CRF receptors 1b (*crfr1b*; C) and 2b (*crfr2b*; D) in the middle intestine (Mid. Int.) of rainbow trout (*Oncorhynchus mykiss*). Significant differences (p < 0.05; one-way ANOVA) are depicted using letters; except in Panel A where the p value is reported because no significant post hoc comparisons were detected. Data are expressed relative to control fish that were fed 1 h prior to sampling. Values are represented as means ± SEM and individual data points are shown (N=6-7).

**Figure 9.**
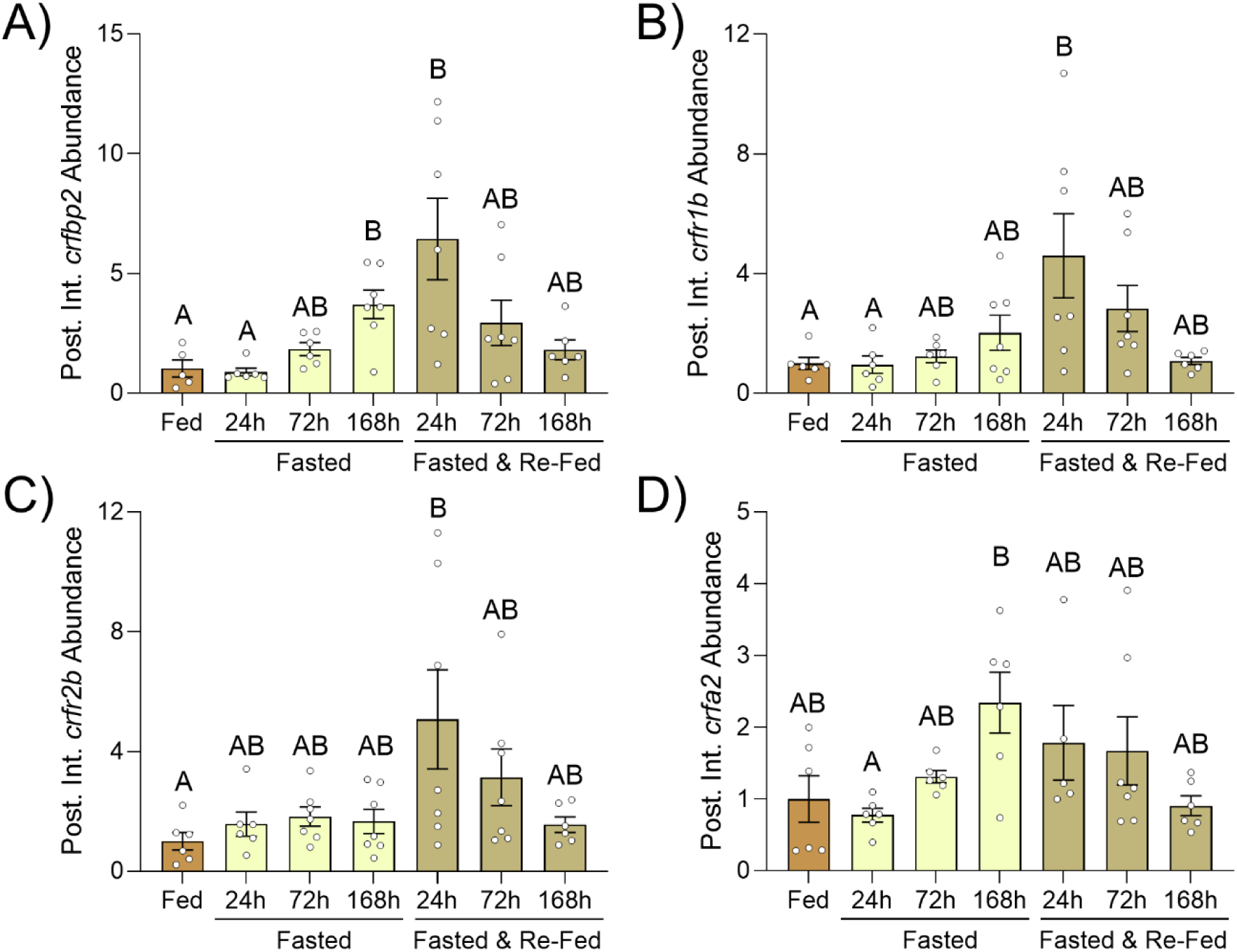
Time-dependent effects of fasting and re-feeding on transcript abundance of corticotropin-releasing factor (CRF) binding protein 2 (*crfbp2*; A), CRF receptors 1b (*crfr1b*; B) and 2b (*crfr2b*; C), and CRFa2 (*crfa2*; D) in the posterior intestine (Post. Int.) of rainbow trout (*Oncorhynchus mykiss*). Significant differences (p < 0.05; one-way ANOVA) are depicted using letters. Data are expressed relative to control fish that were fed 1 h prior to sampling. Values are represented as means ± SEM and individual data points are shown (N=5-7).

While a significant difference in plasma cortisol levels was observed during the fasting-refeeding experiment (Table 1), no differences were detected during subsequent post hoc analysis, and cortisol levels were generally low across all groups (<15 ng mL^-1^).

## 4. Discussion

While neuroendocrine functions for the CRF system in the brain (e.g., regulation of the HPA/HPI axis) are well established (Aguilera, 1998; Best et al., 2024), stress-related roles for the CRF system outside of the brain are less well understood, especially in non-mammalian vertebrates. In the current study, we observed tissue- and stressor-specific transcriptional regulation of the CRF system in the gills and intestine of rainbow trout following different physiological challenges. To determine how the CRF system is affected by an immune challenge, we administered a vaccine and evaluated responses across time. Vaccinated fish displayed lower circulating lymphocytes combined with higher blood neutrophil counts and respiratory burst activity, indicating that fish mounted a robust immune response. Consistent with immune responses observed in the blood, we also detected large (∼15‒300-fold) increases in immune-related transcripts across both the middle and posterior portions of the intestine following vaccination. These responses occurred in parallel with transcriptional changes in several CRF system components 24-72h post-vaccination; specifically, an upregulation of CRFBP (especially *crfbp1*) combined with a downregulation of *crfr1b* and *crfa2*. We also found that splenic levels of these same CRF system components displayed similar transcriptional changes (Culbert et al., 2026, preprint) suggesting that suppression of CRFR1 activity in response to vaccination may be widespread across tissues. In mammals, activation of the CRF system—with specific effects mediated by both CRFR1 and CRFR2—causes wide-ranging proinflammatory effects in the intestine, including increased production of proinflammatory cytokines [e.g., IL6, IL8, and TNFα (Chatoo et al., 2018; Dermitzaki et al., 2018; Kiank et al., 2010)]. While no previous study has evaluated immune contributions of the CRF system in the intestine of fish, the proinflammatory effects triggered by *V. anguillarum* treatment of ayu (*Plecoglossus altivelis*) head kidney leukocytes were attenuated when UTS1 was also added (Jiang et al., 2021), counter to mammalian studies. These anti-inflammatory actions in ayu head kidney leukocytes appear to involve IL6 signalling pathways since UTS1 was able to block the proinflammatory effects caused by exogenous IL6 treatment (Jiang et al., 2021). Consistent with immunosuppressive effects of the CRF system in fish, UTS1 treatment was also associated with reduced phagocytotic activity in cultured spotted snakehead (*Channa punctatus*) spleen monocytes (Singh and Rai, 2011). While our results do not support an anti-inflammatory role for the intestinal CRF system in trout since the observed reduction in CRF system activity coincided with peak production of the anti-inflammatory cytokine *il10*, it is possible that the observed reduction in CRF system activity facilitates greater phagocytotic activity following vaccination. Indeed, Singh & Rai (2011) used receptor-specific antagonists to show that the anti-phagocytotic effects associated with UTS1 were mediated by CRFR1 (and not CRFR2), which is consistent with the observed downregulation of intestinal *crfr1b* at 24h post-vaccination. Additionally, Van Heijningen et al. (2025) reported that mutant zebrafish larvae lacking *crfr1* recruited fewer macrophages and more neutrophils following tail fin amputation, offering further evidence that CRFR1 has important immune functions in teleosts. Overall, while our transcriptional results are consistent with CRFR1-mediated immunoregulatory roles in the intestine of trout, additional work is needed to determine whether these roles vary between species, tissues, cell types, and/or immune components (e.g., innate versus adaptive immune systems).

Unlike in the intestine, we did not detect any transcriptional changes in the gill CRF system following vaccination. As discussed above, since the CRF system generally has proinflammatory effects in mammals, we predicted an acute activation of the CRF system during the initial surge of proinflammatory cytokine activity, followed by suppression of CRF system activity as anti-inflammatory cytokines are activated (Dermitzaki et al., 2018; Webster et al., 1998)—the latter of which was observed in the intestine. Indeed, Jiang et al. (2021) found that transcript levels of UTS1 acutely increased in the gills of ayu following infection with *V. anguillarum* (12-24h after onset of infection). In contrast, Mazon et al. (2006) reported that chronic (3 weeks) infection with the parasite *Trypanoplasma borreli* caused transcript levels of *crfr1* and *crfbp* in the gills of common carp to decrease by ∼60%. These different responses could reflect several factors, including species-specific regulation of the gill CRF system, the type of immune challenge used (vaccination versus infection with an actual pathogen), and/or whether cortisol levels were concurrently elevated. Similarly, it is possible that the immune challenge used in the current study failed to elicit a strong enough local immune response to affect the gill CRF system, especially since previous *in vitro* work has shown that the CRF system can influence proinflammatory cytokine systems in Atlantic salmon (*Salmo salar*) gills [e.g., IL23 and IL17 receptors; (Culbert et al., 2025c)]. While we detected significant transcriptional responses for all immune-related genes that were measured in the gills, these responses were weaker than those observed in the intestine (current study) or spleen (Culbert et al., 2026 preprint), despite similar baseline expression across tissues. However, changes in gill CRF system components were observed by Mazon et al. (2006) despite *T. borreli* infection not being associated with large changes in cytokine expression in carp (Saeij et al., 2003). Therefore, it is possible that changes in the gill CRF system following parasite infection reflect other actions associated with CRF in the gills [e.g., regulation of angiogenesis and/or endothelial permeability; (Culbert et al., 2025c)] and might not have been directly contributing to immunoregulatory functions. Thus, future investigations evaluating how immune challenges of different types and intensities influence the CRF system across tissues and between species will help to determine the extent to which the CRF system serves immune functions in teleosts.

In addition to the immune challenge, we also evaluated how the peripheral CRF system responded to an osmotic stressor. Following SW transfer, fish mounted a cortisol response lasting at least 72h (but not more than 168h) and displayed elevated plasma osmolality during the entire 168h experimental period. Both responses indicate that fish were perturbed by this osmotic stressor (Craig et al., 2005; Culbert et al., 2022); however, the reduction in plasma cortisol levels by 168h suggests that fish had begun to acclimate by this time. Consistent with the plasma responses observed, we also detected transcriptional changes in several components of the gill CRF system during this period. Most notably, *crfr1a* and *crfr2a* were downregulated across all sampling times in SW transferred fish, while levels of *crfr1b* and *crfbp1* were lower in SW transferred fish at 72 and 168h, respectively. These findings are largely consistent with our previous work in Atlantic salmon, where SW transfer was associated with a general transcriptional downregulation of the CRF system (and vice-versa when SW-acclimated salmon were transferred into FW)—especially *crfr2a* and *ucn2a* (Culbert et al., 2025c). Indeed, the tissue distribution of CRF system components in rainbow trout (current study) and Atlantic salmon (Culbert et al., 2025c) also shares many similarities, especially high expression of *crfr2a* and *crfbp1*, suggesting that the role(s) of the CRF system in the gills are likely conserved among salmonids. We have previously suggested that these changes in gill CRF system transcriptional activity reflect regulatory contributions of the CRF system towards angiogenesis, endothelial permeability, and/or immune regulation in the gills because *in vitro* treatment of gill filaments with CRFa affected levels of transcripts related to these physiological processes (Culbert et al., 2025c). However, it is likely that the role of the gill CRF system varies between species since levels of *crfr1* and *crfb* were ∼5-15-fold higher in tilapia (*Oreochromis mossambicus*) 30 days after transfer from FW-to-SW (Aruna et al., 2012), contrary to the downregulation of CRFR1 observed here. Our previous work in Atlantic salmon (Culbert et al., 2025c) also indicates that the observed downregulation of *crfr1a* and *crfr1b* (at least during the first 72h following SW transfer) could be mediated by elevated cortisol levels. Yet, there also appear to be species-specific differences in the regulatory effects of corticosteroids on components of the gill CRF system since *crfr1* levels were increased when black porgy (*Acanthopagrus schlegelii*) gill filaments were treated *in vitro* with dexamethasone for 48 h (Aruna et al., 2021). Regardless, the current data clearly support the hypothesis that the CRF system has osmoregulatory functions in the gills of salmonids.

In contrast to the gills, while transcript levels of several CRF system components in the Mid. Int. were initially downregulated following SW transfer (*crfbp2*, *crfr1b*, and *crfr2b*), the abundance of most CRF system components was elevated by 168h post-transfer to SW. These contrasting responses between tissues likely reflect the different osmoregulatory role(s) of the gills versus the intestine in SW. Whereas the gills are responsible for exporting ions out of the body, ions are taken up across the intestine to maintain a favourable gradient facilitating water absorption (Evans et al., 2005; Grosell, 2010; Larsen et al., 2014). However, it is not clear why greater CRF system activity in the intestine of SW-acclimated fish would be beneficial from an osmoregulatory standpoint since the CRF system generally promotes ion and/or water excretion (or reduces absorption) across taxa and epithelium types (Cannell et al., 2016; Marshall and Bern, 1981; Stengel and Taché, 2009). Indeed, activation of CRFR2 in the intestine of FW-acclimated Atlantic salmon reduced absorption of water, Na^+^, and Cl^-^ (Culbert et al., 2025b). Similarly, Mainoya & Bern (1982) reported that UTS1 treatment had inhibitory effects on water, Na^+^, and Cl^-^ uptake in the intestine of FW-acclimated (but not SW-acclimated) Mozambique tilapia, suggesting a greater role for the intestinal CRF system in FW. Therefore, it is possible that the observed changes in the CRF system during SW acclimation may indirectly assist with water and/or solute transport by affecting processes related to angiogenesis and/or endothelial permeability in the Mid. Int. (Culbert et al., 2025c). Unlike the Mid. Int., few changes in CRF system transcript levels were observed in the Post. Int. following SW transfer. Instead, transcript levels of several genes (*crfbp1*, *crfbp2*, *crfr1b*, *crfa2*, and *ucn2b*) increased across time in both the FW- and SW-transferred groups. Indeed, time-dependent increases in *crfa2* transcript levels were also observed in the Post. Int. during the vaccination experiment. Since fish were fasted during both experiments—replicating natural responses of salmonids following immune and osmotic stressors (Arnesen et al., 1993; Craig et al., 2005; Damsgård and Arnesen, 1998; Sørum and Damsgård, 2004; Usher et al., 1991)—we decided to evaluate whether changes in feeding state might explain these responses.

The hypothalamic CRF system is a potent regulator of food intake in fish and other vertebrates (Bernier, 2006; Conde-Sieira et al., 2018; Stengel and Taché, 2014), but the relationship between food intake and CRF system components located outside of the brain is less clear. Several previous studies have reported the presence of various CRF system components in the teleost intestine (Arai et al., 2001; Chen and Fernald, 2008), but we believe that our data are the first to show that transcript abundance of CRF system components in the intestine can vary with feeding state. Interestingly, responses in the middle versus posterior regions of the intestine exhibited opposing patterns, wherein levels declined in the Mid. Int. and increased in the Post. Int. As discussed earlier, activation of the CRF system reduces absorption of water and ions across the intestine in fish and the colon in mammals (Culbert et al., 2025b; Mainoya and Bern, 1982; Santos et al., 1999). While these effects are mediated by CRFR1 in mammals (Stengel and Taché, 2009), activation of CRFR2 was primarily responsible for these effects in Atlantic salmon (Culbert et al., 2025b). Interestingly, peripheral treatment with CRF peptides can also delay clearance rates in the stomach and small intestine via a CRFR2-dependent mechanism in mammals (Stengel and Taché, 2009). Thus, changes in CRF system activity observed in both the middle (reduced abundance, suggesting faster clearance rates) and posterior (increased abundance, suggesting reduced absorption rates) regions of the intestine are consistent with CRF contributing to reductions in intestine functionality and investment during extended fasts. While the functional importance of such CRF-mediated changes is unclear, they may contribute to fasting-related changes in nutrient/water absorption, immune regulation, and/or alterations in the microbiome (Li et al., 2017; Wood and Bucking, 2010; Xia et al., 2014). However, since fasting-related transcriptional responses in both regions of the intestine were generally different than those following either SW transfer or vaccination, the role of the intestinal CRF system is clearly complex and warrants additional study.

In conclusion, we found that the gill and intestine CRF systems are differentially regulated by osmotic and immune stressors. While activity of the gill CRF system is reduced following SW transfer, we found no evidence of a role for the gill CRF system during an immune stressor. In contrast, vaccination was associated with reduced activity of the intestine CRF system, and SW transfer transcriptionally activated the CRF system in the Mid. Int. Lastly, fasting caused region-specific changes in CRF system activity within the middle (reduction) and posterior (activation) regions of the intestine. Overall, our findings highlight the breadth and functional diversity of stress-related roles that the CRF system serves outside of the brain.

## Funding

This work was supported by a Natural Sciences and Engineering Research Council of Canada (NSERC) Discovery grant provided to NB (RGPIN-2022-03151). BMC was supported by a NSERC Doctoral Canadian Graduate Scholarship (CGS-D) and an Ontario Graduate Scholarship (OGS). CB was supported by a NSERC postdoctoral scholarship.

## Acknowledgments

We thank Matt Cornish, Carolyn Trombley, and Mike Davies for their assistance with fish care and advice regarding experimental setups, as well as Dr. Marcia Chiasson and the staff at the Ontario Aquaculture Research Centre for providing us with the rainbow trout. We also thank Emma Mossington and Shayla Larson for their help during dissections, Sarah Alderman for allowing use of the flow cytometer, and Brian Dixon and Tania Rodríguez-Ramos for assistance with the respiratory burst assay.

## Data accessibility

Data are deposited on Mendeley Data (doi: 10.17632/bnr9zp28fp.1).

## Notes

https://data.mendeley.com/datasets/bnr9zp28fp/1

### Competing Interest Statement

The authors have declared no competing interest.

