## Supplemental Tables for "Stressor- and tissue-specific regulation of the corticotropin-releasing factor system across epithelial tissues in rainbow trout"

Brett M. Culbert^*†^, Alexis E. Pulford-Thorpe, Carol Best,

& Nicholas J. Bernier

*Department of Integrative Biology, University of Guelph, Guelph, Ontario, Canada*

^†^Current Address: *Department of Biological Sciences, University of Cincinnati, Cincinnati, Ohio, USA*

**Supplemental Table 1.** Gene specific primers used for real-time qPCR.

|  | Primer Sequence  (5’ to 3’) | Amplicon  Size (bp) | Efficiency  (%) | Accession  Number |
| --- | --- | --- | --- | --- |
| *crfa1* | F: CAAAATTCGCTCCAGTCCTC | 95 | 100 | XM_021613200 |
|  | R: TGCGTGAGCTGAAGTTGTAA |  |  |  |
| *crfa2* | F: ATTCGCCCCAATCTTCATT | 79 | 102 | XM_021589550 |
|  | R: TGAAGTAAAGCCCTGTTGACC |  |  |  |
| *crfb1* | F: CTGAAGGCATGTACCCAGAG | 110 | 102 | NM_001124286 |
|  | R: GACGAACCGGCTGATGTT |  |  |  |
| *crfb2* | F: GAAAGTCATGCACCCCAAG | 117 | 102 | NM_001124627 |
|  | R: AAGCCGCTGATGAACCTATT |  |  |  |
| *crfbp1* | F: CCAACACGTTGTCTATCTAC | 107 | 100 | NM_001124631 |
|  | R: TTTTATGGCAACCTTCAGGCT |  |  |  |
| *crfbp2* | F: CTCCCTATCTATCTCTCCAC | 122 | 100 | XM_036977080 |
|  | R: GTTATGGCGATCCTCAGGAT |  |  |  |
| *crfr1a* | F: TCCCTCGGAGACAGGAGTGT | 121 | 102 | XM_036940770 |
|  | R: CTTTTACGATGTGCGACTGGT |  |  |  |
| *crfr1b* | F: CGGCAATTTATAGGCTGTGTT | 114 | 99 | XM_021624572 |
|  | R: CCCACATGAACAACCAACAG |  |  |  |
| *crfr2a* | F: CACTGTACTGTACTAGTGGT | 83 | 100 | XM_021613636 |
|  | R: TCCTGCTTAAATGATCTCTGC |  |  |  |
| *crfr2b* | F: TAAATTCAGAACAGATGGGC | 82 | 100 | XM_021589160 |
|  | R: CATTGCCAGTAAGACTCTGAC |  |  |  |
| *c3* | F: GAGCAGCACCTGGCTGACA | 150 | 102 | XM_021561577 |
|  | R: GGAGCAAACTCATTGAAGATGCC |  |  |  |
| *ef1α* | F: CCATTGACATTTCTCTGTGGAAGT | 106 | 96 | NM_001124339 |
|  | R: GAGGTACCAGTGATCATGTTCTTGA |  |  |  |
| *ifnγ* | F: ACCCTGTTTTCCCCAAGGAC | 61 | 92 | NM_001124620 |
|  | R: ACCACACTCATCAACACCCTC |  |  |  |
| *il1b* | F: TGAGAACAAGTGCTGGGTCC | 148 | 107 | NM_001124347 |
|  | R: GGCTACAGGTCTGGCTTCAG |  |  |  |
| *il10* | F: CCGCCATGAACAACAGAACA | 105 | 93 | NM_001245099 |
|  | R: TCCTGCATTGGACGATCTCT |  |  |  |
| *rpl13a* | F: GGACAAGCTGCACTGGAGAG | 113 | 100 | XM_021574753 |
|  | R: GTGGGCTTCAGACGGACAAT |  |  |  |
| *tnfα* | F: CACACTGGGCTCTTCTTCGT | 155 | 98 | NM_001124374 |
|  | R: CAAACTGACCTTACCCCGCT |  |  | NM_001124357 |
| *ucn2a* | F: CATGCGTATGTGTCTGAGCC | 61 | 101 | CDQ61544 |
|  | R: TCTTCTCCCATCCTTTGGCAC |  |  |  |
| *ucn2b* | F: ACTCCTAAGTTGTGACATTGGA | 68 | 99 | XM_021570508 |
|  | R: CCAGACACAGGCTTAGATG |  |  |  |
| *ucn3* | F: GCACGGAACTTGGACATCAT | 91 | 100 | XM_021596023 |
|  | R: GTCTGAGACAGAGGCTGGAG |  |  | XM_036945073 |
| *uts1a* | F: GTTGCTGAAGGCTGGTGATA | 82 | 100 | NM_001124343 |
|  | R: CAACCGGGGATTTCTTTG |  |  |  |
| *uts1b* | F: GGCGACGCAGTTTCCTAC | 60 | 99 | XM_021612039 |
|  | R: GGGATTTCTCTGCAAATAACG |  |  |  |

*crf*, corticotropin-releasing factor; *crfbp*, corticotropin-releasing factor binding protein; *crfr*, corticotropin-releasing factor receptor; *c3*, complement 3; *ef1α*, elongation factor 1α; *ifnγ* interferon γ, *il*, interleukin; *rpl13a*, ribosomal protein L13a; *tnfα*, tumor necrotic factor α; *ucn*, urocortin; *uts*, urotensin.

**Supplemental Table 2.** Time dependent effects of saline or vaccine treatment on transcript abundance of CRF system components in the gills, and middle (Mid. Int.) and posterior (Post. Int.) segments of the intestine in rainbow trout (*Oncorhynchus mykiss*). Data are presented as means ± SEM (N = 8=11 per group). Significant effects are indicated with **bold** (p < 0.05) and differences based on post hoc analysis are indicated using letters (across time within a group), asterisks (between groups within a timepoint) or symbols (overall time effects).

|  |  |  | Sampling Time | | | |  |  |  |
| --- | --- | --- | --- | --- | --- | --- | --- | --- | --- |
| Tissue | Gene | Treatment | 6 h | 24 h | 72 h | 168 h |  | F | p |
| Gills | *crfbp1* | Saline | 1.00±0.14^α^ | 1.04±0.07^αβ^ | 1.02±0.11^αβ^ | 1.22±0.09^β^ | Treatment | 0.3 | 0.6 |
|  |  | Vaccine | 0.84±0.08^α^ | 0.99±0.07^αβ^ | 1.09±0.09^αβ^ | 1.21±0.09^β^ | **Time** | **3.4** | **0.02** |
|  |  |  |  |  |  |  | Interaction | 0.5 | 0.67 |
|  | *crfr1a* | Saline | 1.00±0.10 | 1.02±0.09 | 1.00±0.07 | 0.88±0.11 | Treatment | 0.1 | 0.71 |
|  |  | Vaccine | 0.92±0.11 | 0.96±0.10 | 0.86±0.10 | 1.10±0.13 | Time | 0.2 | 0.92 |
|  |  |  |  |  |  |  | Interaction | 1.4 | 0.27 |
|  | *crfr1b* | Saline | 1.00±0.06 | 0.91±0.08 | 0.77±0.08 | 0.99±0.10 | Treatment | 0.7 | 0.40 |
|  |  | Vaccine | 0.94±0.08 | 0.85±0.10 | 0.80±0.07 | 0.87±0.08 | Time | 2.1 | 0.11 |
|  |  |  |  |  |  |  | Interaction | 0.2 | 0.88 |
|  | *crfr2a* | Saline | 1.00±0.06^A^ | 0.69±0.04^B^ | 0.48±0.05^C*^ | 0.72±0.06^A^ | **Treatment** | **4.5** | **0.04** |
|  |  | Vaccine | 0.86±0.07 | 0.87±0.10 | 0.72±0.04 | 0.76±0.05 | **Time** | **10.6** | **<0.001** |
|  |  |  |  |  |  |  | **Interaction** | **4.2** | **0.009** |
|  | *crfr2b* | Saline | 1.00±0.08 | 1.06±0.09 | 0.93±0.10 | 0.82±0.07 | Treatment | 0.1 | 0.71 |
|  |  | Vaccine | 0.90±0.07 | 1.03±0.08 | 0.79±0.05 | 1.21±0.14 | Time | 1.5 | 0.22 |
|  |  |  |  |  |  |  | Interaction | 2.6 | 0.06 |
|  | *ucn3* | Saline | 1.00±0.12 | 1.06±0.16 | 2.22±0.72 | 3.06±0.82 | Treatment | 1.8 | 0.18 |
|  |  | Vaccine | 1.53±0.28 | 0.70±0.09 | 1.27±0.31 | 0.93±0.08 | Time | 1.4 | 0.25 |
|  |  |  |  |  |  |  | Interaction | 1.3 | 0.27 |
| Mid.  Int. | *crfr2b* | Saline | 1.00±0.21^αβ^ | 0.89±0.29^α^ | 1.38±0.11^β^ | 1.20±0.16^αβ^ | Treatment | 0.6 | 0.44 |
|  |  | Vaccine | 1.05±0.17^αβ^ | 0.56±0.14^α^ | 1.40±0.16^β^ | 1.07±0.10^αβ^ | **Time** | **4.9** | **0.004** |
|  |  |  |  |  |  |  | Interaction | 0.5 | 0.68 |
|  | *ucn2b* | Saline | 1.00±0.20 | 1.07±0.26 | 1.72±0.21 | 1.40±0.19 | Treatment | 1.1 | 0.29 |
|  |  | Vaccine | 1.21±0.08 | 1.74±0.33 | 1.09±0.20 | 1.92±0.35 | Time | 1.4 | 0.24 |
|  |  |  |  |  |  |  | Interaction | 2.7 | 0.05 |
| Post.  Int. | *crfbp2* | Saline | 1.00±0.24 | 1.06±0.20 | 1.53±0.25 | 1.27±0.25 | Treatment | 0.1 | 0.85 |
|  |  | Vaccine | 0.74±0.14 | 1.54±0.34 | 1.30±0.22 | 1.06±0.16 | Time | 2.1 | 0.11 |
|  |  |  |  |  |  |  | Interaction | 0.7 | 0.57 |
|  | *crfr2b* | Saline | 1.00±0.33 | 1.12±0.30 | 0.90±0.09 | 0.94±0.20 | Treatment | 0.6 | 0.45 |
|  |  | Vaccine | 0.95±0.23 | 0.81±0.12 | 1.01±0.12 | 0.91±0.10 | Time | 1.0 | 0.41 |
|  |  |  |  |  |  |  | Interaction | 0.2 | 0.93 |
|  | *ucn2b* | Saline | 1.00±0.20 | 1.19±0.24 | 0.87±0.17 | 1.00±0.16 | Treatment | 1.0 | 0.33 |
|  |  | Vaccine | 1.05±0.20 | 1.35±0.28 | 0.97±0.12 | 1.12±0.18 | Time | 0.8 | 0.51 |
|  |  |  |  |  |  |  | Interaction | 0.1 | 0.99 |

**Supplemental Table 3.** Time-dependent effects of transfer from freshwater-to-freshwater (FW) or freshwater-to-seawater (SW) on transcript abundance of CRF system components in the gills and posterior intestine (Post. Int.) in rainbow trout (*Oncorhynchus mykiss*). Data are presented as means ± SEM (N = 6=10 per group).

|  |  |  | Sampling Time | | |  | |  | |  | |
| --- | --- | --- | --- | --- | --- | --- | --- | --- | --- | --- | --- |
| Tissue | Gene | Treatment | 24 h | 72 h | 168 h | |  | | F | | p |
| Gills | *crfr2b* | FW | 1.00±0.13 | 1.24±0.21 | 0.88±0.08 | | Treatment | | 0.1 | | 0.87 |
|  |  | SW | 0.83±0.10 | 0.99±0.11 | 1.24±0.16 | | Time | | 1.3 | | 0.28 |
|  |  |  |  |  |  | | Interaction | | 3.1 | | 0.06 |
|  | *ucn3* | FW | 1.00±0.26 | 0.84±0.20 | 1.34±0.33 | | Treatment | | 3.2 | | 0.08 |
|  |  | SW | 0.89±0.34 | 0.48±0.12 | 0.82±0.22 | | Time | | 1.2 | | 0.30 |
|  |  |  |  |  |  | | Interaction | | 0.1 | | 0.92 |
| Post.  Int. | *crfr2b* | FW | 1.00±0.15 | 1.49±0.48 | 1.01±0.18 | | Treatment | | 1.4 | | 0.24 |
|  |  | SW | 1.27±0.26 | 1.33±0.28 | 1.77±0.33 | | Time | | 0.6 | | 0.57 |
|  |  |  |  |  |  | | Interaction | | 1.0 | | 0.39 |

**Supplemental Table 4.** Time-dependent effects of fasting and refeeding on transcript abundance of CRF system components in the middle (Mid. Int.) and posterior (Post. Int.) segments of the intestine in rainbow trout (*Oncorhynchus mykiss*). Data are presented as means ± SEM (N = 6-7 per group). Significant effects are indicated with **bold** (p < 0.05) and differences between groups based on post hoc analysis are indicated using letters. Note that significant differences were not detected during post hoc analysis of *crfbp1* in the Mid. Int.

|  |  |  | Fasting | | |  | Refeeding | | |  |  |
| --- | --- | --- | --- | --- | --- | --- | --- | --- | --- | --- | --- |
| Tissue | Variable | Fed | 24 h | 72 h | 168 h |  | 24 h | 72 h | 168 h | F | p |
| Mid.  Int. | *crfa2* | 1.00±0.17 | 1.40±0.41 | 0.75±0.05 | 2.01±0.57 |  | 0.95±0.13 | 0.82±0.13 | 0.79±0.15 | 2.2 | 0.06 |
|  | *ucn2b* | 1.00±0.16 | 0.36±0.03 | 0.65±0.17 | 0.71±0.19 |  | 0.59±0.14 | 0.53±0.10 | 0.99±0.28 | 1.3 | 0.26 |
| Post.  Int. | *ucn2b* | 1.00±0.33 | 0.65±0.14 | 0.91±0.25 | 0.95±0.15 |  | 1.18±0.23 | 0.86±0.15 | 0.64±0.13 | 0.7 | 0.65 |
|  | *crfbp1* | 1.00±0.26 | 1.57±0.48 | 0.88±0.12 | 0.81±0.18 |  | 1.15±0.22 | 0.77±0.09 | 1.13±0.36 | 1.1 | 0.40 |
